# Synergy and convergence of pathways controlling functional regeneration in the spinal cord

**DOI:** 10.1101/334029

**Authors:** Lucas P. Carlstrom, Thomas R. Cheever, Heiko L. Schoenfuss, Meghan R. McGee, Stephen C. Ekker, John R. Henley

## Abstract

Barriers to regeneration in the mammalian central nervous system (CNS) include the presence of inhibitory factors like myelin-associated glycoprotein (MAG) that block re-growth of injured axons. Inhibition by MAG antagonizes the induction of integrin-based substrate adhesions in axonal growth cones by brain-derived neurotrophic factor (BDNF). Here, using a novel approach to overcome inhibitory actions of MAG by activating integrins, we provide cellular and molecular evidence that integrin activity modulates the actions of chemotropic cues on substrate adhesions and supports axon regeneration in vertebrates. Potentiating integrin activity in cultured spinal neurons blocked negative integrin remodeling and inhibition of axon outgrowth induced by MAG, but also restored BDNF-dependent integrin clustering and stimulated outgrowth. In a zebrafish complete spinal cord transection model, combined integrin activation and BDNF treatment synergistically triggered functional regeneration of long projection axons that lack regenerative capacity from the hindbrain. The combined treatment also promoted functional repair even in the presence of exogenous mammalian inhibitory factors, including MAG, which alone impaired recovery of swimming movements. Thus, integrin activation state plays complementary roles in modulating the output activity of opposing cues on integrin-based adhesions and supports functional nerve regeneration *in vivo*. Our findings reveal effective reversal of downstream actions of inhibitory cues, thereby overcoming a major barrier to regeneration in the mammalian CNS, while simultaneously supporting neurotrophin-stimulated outgrowth. Discovery of therapeutic strategies targeting integrin activation state therefore holds promise for promoting axon regeneration after traumatic injury, which is a critical step in restoring connectivity and functional recovery.

## INTRODUCTION

Features of the injured mammalian spinal cord that prevent meaningful natural regeneration include diminished intrinsic axon growth capacity and a complex, growth-prohibitive environment^1,2^. The non-permissive injury milieu contains not only factors released from myelin debris, like MAG^3-5^, but also chondroitin sulfate proteoglycans (CSPGs) and semaphorin 3A (Sema 3A), which are known potent inhibitors of axon outgrowth^1^. Thus, design of an effective therapy to induce functional regeneration likely will need to overcome the effects of these inhibitory factors in addition to stimulating axon growth capacity. Understanding the mechanisms underlying both inhibition of axon outgrowth and neurotrophin-dependent stimulation of outgrowth may provide critical insights towards this goal.

Previous attempts to block the actions of inhibitory cues have had limited success, indicating an incomplete understanding of the underlying mechanisms. Emerging evidence suggests that multiple cues affecting axon outgrowth may converge on a common downstream cellular process regulating integrin adhesion complexes^6-8^. Our previous work demonstrated that solubilized MAG strongly promotes endocytic removal of β1-integrin from the nerve growth cone plasmalemma, which is both necessary and sufficient to drive chemorepulsion^6^. Furthermore, CSPGs and Sema 3A impair the activation of β1-integrin and lead to de-aggregation of adhesion proteins in the growth cone^9-13^, providing evidence that negative regulation of integrin adhesions is a common downstream mechanism of inhibitory molecules. In contrast, BDNF positively regulates integrin clustering and the formation of adhesions during axon attraction and stimulated outgrowth^14,15^. Interestingly, overexpression of integrins can promote neurite growth in culture on weak substrates^16,17^, and treatment with Mn^2+^ to activate integrins can partially overcome inhibition of axon outgrowth by Nogo^7,11^ and CSPGs in cell culture^11^. Moreover, co-upregulating expression of a weak substrate-binding integrin and the integrin activating protein kindlin-1 stimulates regenerative growth of sensory axons after dorsal root crush injury^18^. Thus, the individual actions of axon outgrowth inhibiting and stimulating factors intersect on a common cellular target that may be critical for regeneration^16-19^. We here explore the role of integrin activity in supporting neurotrophin-dependent axon outgrowth and neural regeneration in the presence of mammalian inhibitory cues.

## RESULTS

### Activating integrins restores BDNF-induced adhesions and axon outgrowth

Whereas the cellular actions of MAG and BDNF converge on integrin receptors and axon outgrowth, how potentiating integrin activity might modulate these interactions remains unexplored. We therefore investigated how inducing integrins into a conformationally activated state^20^ affects the functional distribution of integrin receptors and axon outgrowth during exposure to these opposing factors. In primary *Xenopus laevis* spinal neurons, exposure to MAG disrupts β1-integrin clustering at the growth cone, reduces surface levels, and impairs rates of axon outgrowth^6,14^. Treatment with Mn^2+^, which potentiates integrin activity, blocks integrin internalization by MAG, and increases β1-integrin surface levels slightly^6^, was permissive for stimulation of integrin clustering and axon outgrowth by BDNF (Fig. 1a-d). Moreover, pretreatment with Mn^2+^ followed by exposure to MAG and then BDNF administration restored integrin clustering and stimulated axon outgrowth (Fig. 1a-d). Intriguingly, initial MAG exposure followed by post-treatment with Mn^2+^ and BDNF together was sufficient to induce integrin adhesion formation and stimulated axon extension (Fig. 1a-d). This is in contrast to the loss of integrin adhesions, reduced surface levels, and impaired axon outgrowth induced by initial MAG exposure and followed by BDNF treatment alone. Initial MAG exposure followed by addition of a β1-integrin activating antibody and BDNF combined also led to a substantial restoration of axon outgrowth similar to the combined treatment with Mn^2+^ (Supplementary Fig. S1). Thus, combinatorial treatments to target integrin avidity can partially reverse the negative remodeling of integrin adhesions induced by MAG while permitting the stimulatory effects of a pro-growth factor.

**Figure 1.**
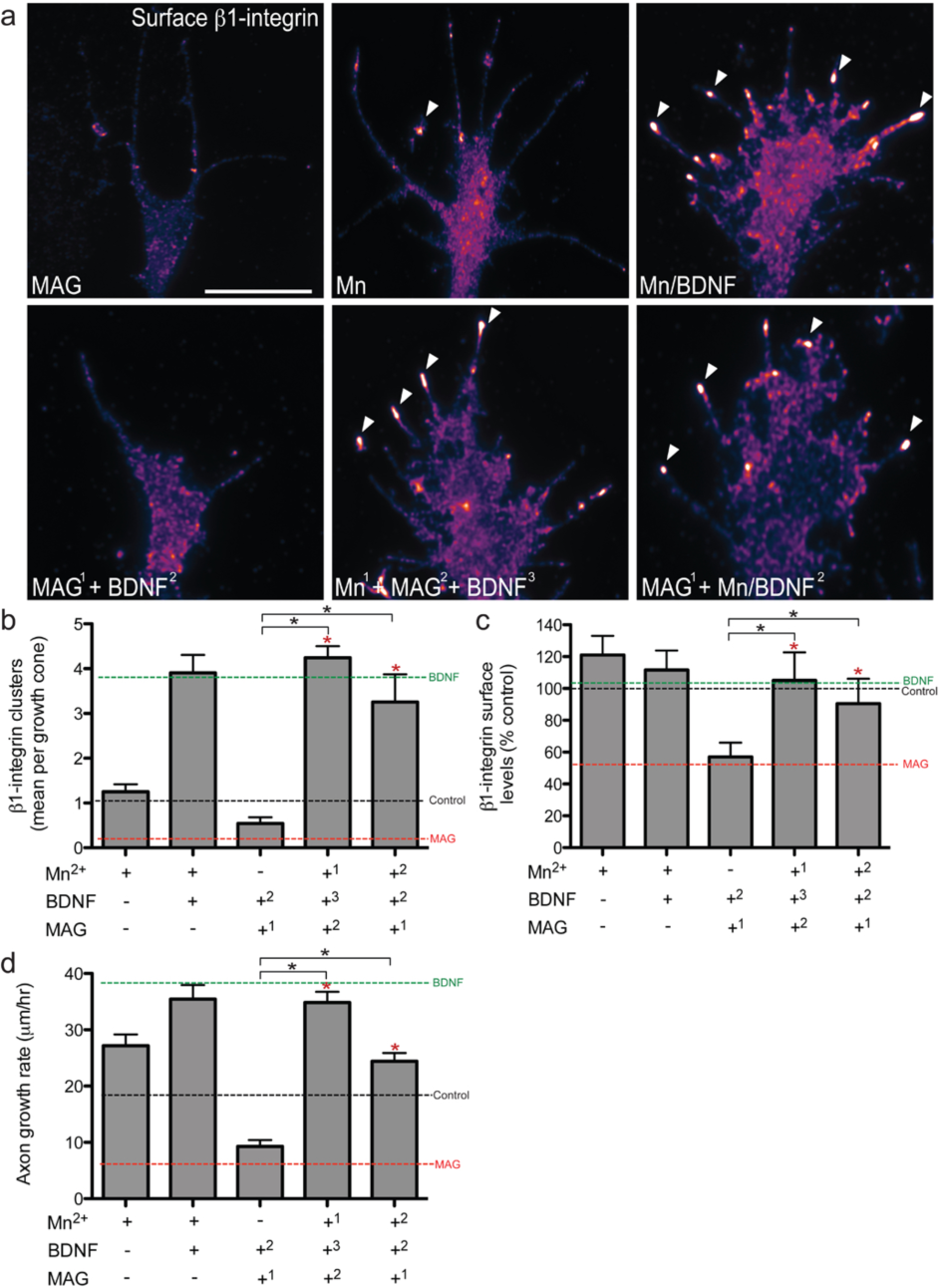
Integrin activation blocks the negative effects of MAG and restores BDNF-induced formation of integrin adhesions and stimulated axon outgrowth. *a*, Representative images show β1-integrin immunolabeling in *Xenopus* axonal growth cones after MAG (1 µg mL^−1^; 5 min), Mn^2+^ (1 mM; 20 min) or combination treatments with Mn^2+^ and BDNF (50 ng mL^−1^; 20 min), MAG then BDNF, Mn^2+^ followed by MAG and BDNF, and MAG followed by Mn^2+^ and BDNF. Superscript numbers indicate order of treatments applied. Arrowheads designate clustered β1-integrin. Scale bar, 5 µm. Quantification of β1-integrin clustering (*b*), β1-integrin surface levels (*c*) and live rates of axon outgrowth (*d*). Colored dash lines represent the mean of individual application of BDNF (green), control (BSA; black) or MAG (red). Data are the mean ± standard error of the mean. (n > 100, *P < 0.01, **P < 0.001, ANOVA with Tukey’s post hoc analysis). BDNF: brain-derived neurotrophic factor; BSA: bovine serum albumin; MAG: myelin-associated glycoprotein.

### Combined Mn^2+^ and BDNF treatment enhances natural growth capacity

To determine whether using integrin-activating agents can stimulate regenerative growth and functional recovery *in vivo*, we utilized a novel larval zebrafish (*Danio rerio*) complete spinal cord transection model (Supplementary Fig. S2). The zebrafish makes for an attractive model to study regeneration in the spinal cord due to its optical clarity and because it contains specific neurons that fail to regenerate as well as a subset of neurons that can^21-23^. Larval zebrafish are also thought to maintain a largely growth-permissive environment at the injury site that lacks many of the impediments found in mammals^22-26^. Zebrafish have two β1-integrin co-orthologs that have high sequence similarity to the single gene found in humans, including >89% and >99% sequence identity for the ligand binding domain and transmembrane-cytoplasmic region, respectively^27^. Both orthologs are highly expressed in zebrafish larvae and in the adult CNS^27^. Activated β1-integrin receptors are present at the growth cone plasmalemma of cultured zebrafish spinal neurons, and this receptor activation is augmented by Mn^2+^ treatment (Supplementary Fig. S3).

Mauthner neurons (M-cells) are a pair of hindbrain reticulospinal neurons with long myelinated axons that project longitudinally the length of the spinal cord. The M-cells serve as command neurons for the evoked predator escape behavior termed the “C-start”^28^ (Supplementary Movie S1). Despite considerable natural regeneration in the zebrafish spinal cord, M-cell axons demonstrate negligible re-growth after transection (Fig. 2a,b,d,e), as noted in previous reports^22,24^. Remarkably, treatment with Mn^2+^ and BDNF combined for 3 d post-injury (dpf) revealed robust M-cell axon extension caudally with a median of 1727 mm ± 549 beyond the lesion site compared to 238 mm ± 138 for the control-treated (BSA) injured group (Fig. 2b-e). This stands in contrast to fish treated with BDNF alone, which showed only a partial regenerative growth of the M-cell axon when compared to control-treated injured fish (Supplementary Fig. S4). Altogether, this work demonstrates the effectiveness of how targeting integrin activity and neurotrophin signaling together can promote axon regenerative growth *in vivo* of CNS neurons that fail to naturally regenerate after injury.

**Figure 2.**
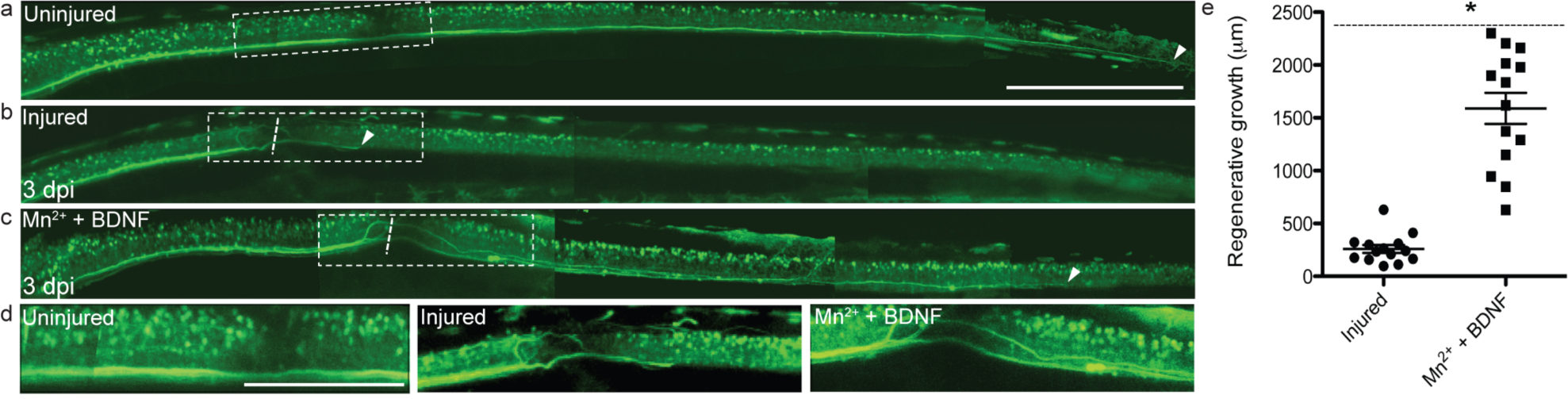
Combination Mn^2+^ and BDNF treatment promotes regenerative axon growth after complete spinal cord transection in larval zebrafish. Multi-photon imaging of the M-cell axon in the transgenic zebrafish line Tg(J1229a:GFP) of uninjured control-treated (BSA) (*a*), injured control-treated (BSA) (*b*), and injured Mn^2+^ (1 mM) + BDNF (100 ng mL^−1^) treated zebrafish 1 h after injury (*c*). Dashed boxes indicate the region of enhanced zoom. White arrowheads denote the most distal region of the M-cell axon detected. All treatments were bath applied. Scale bar, 500 µm. *d*, Expanded detail of the white dashed line box from *a-c* of the lesion site. *e*, Quantitation of regenerative growth observed from injured control-treated (BSA) and injured Mn^2+^ + BDNF treated zebrafish. Dashed line indicates the mean length of the uninjured M-cell axon from the lesion site. Data are the mean ± standard error of the mean. (*P<0.01, ANOVA with Tukey’s post hoc analysis). BDNF: brain-derived neurotrophic factor; BSA: bovine serum albumin.

### Functional recovery through targeting integrin receptors

To test whether the regenerative axon growth from combined Mn^2+^ and BDNF treatment resulted in functional recovery, we evaluated restoration of the larval zebrafish C-start. In response to an external vibration stimulus, uninjured zebrafish rapidly bend into a “C” shape (Fig. 3a; Supplementary Movie S1). Transection of M-cell axons strongly attenuates this behavior, and results in minimal bending along the body axis (Fig. 3a; Supplementary Fig. S5; Movie S2). Remarkably, fish that were co-administered Mn^2+^ and BDNF post injury showed near complete recovery of the C-start response (Fig. 3a-c; Supplementary Fig. S5; Movie S3). Individual treatments with either Mn^2+^ or BDNF alone showed only partial recovery (Fig. 3b,c; Supplementary Fig. S5). In addition to restoration of this evoked behavior, quantitative evaluation of spontaneous swimming activity showed a sharp decline after injury (Fig. 3d,e). This reduction in swimming behavior was reversed with Mn^2+^ and BDNF administration combined (Fig. 3d,e). The restoration of swimming behavior was also observed with a combined β1-integrin activating antibody and BDNF treatment (Fig. 3e). In contrast, pre-treatment with a β1-integrin function-blocking antibody abolished the functional regeneration from Mn^2+^ and BDNF administration (Fig. 3e). Additionally, downregulating expression of both β1-integrin co-orthologs by targeted delivery of antisense morpholino oligonucleotides^27^ (Supplementary Fig. S3) to injured axons strongly attenuated spontaneous swimming activity and abolished the positive functional recovery seen from Mn^2+^ and BDNF treatment (Supplementary Fig. S6). Taken together, these results demonstrate that integrin activating agents combined with neurotrophin treatment can act synergistically to stimulate β1-integrin dependent functional axon regeneration *in vivo*.

**Figure 3.**
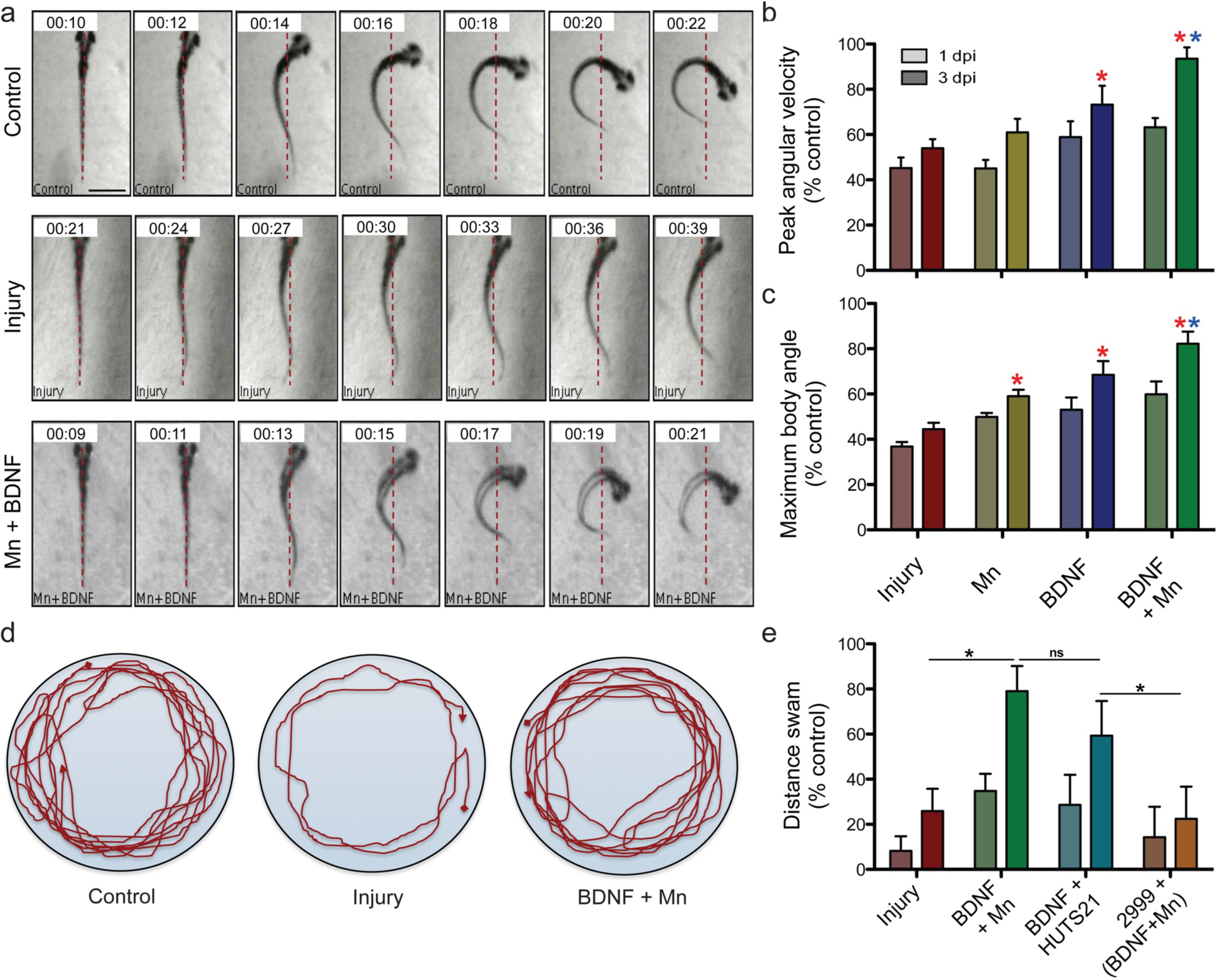
Functional recovery through targeting integrin receptors. *a*, Evoked escape response “C-start” assay depicting high-speed video capture (1,000 Hz) of larval zebrafish immediately preceding response to a vibration stimulus and ending with maximal body angle achieved for uninjured control-treated (BSA), injured control-treated (BSA), and injured Mn^2+^ (1 mM) + BDNF (100 ng mL^−1^) treated zebrafish. Scale bar, 1 mm. *b,c*, Quantification of evoked escape response for control treated (BSA), Mn^2+^ treated, BDNF treated, and Mn^2+^ + BDNF treated injured zebrafish as a percentage of uninjured control-treated (BSA) zebrafish at 1 and 3 d after injury for peak angular velocity (*b*) and maximum body angle (*c*). *d*, Representative tracing of spontaneous zebrafish swimming behavior for uninjured control-treated (BSA), injured control-treated (BSA), and injured Mn^2+^ + BDNF treated zebrafish (6 dpf, 3 dpi for injured fish). *e*, Quantitative MATLAB analysis of spontaneous swimming activity of larval zebrafish at 1 and 3 d after complete spinal cord transection for control-treated (BSA), Mn^2+^ + BDNF treated, β1- integrin activating antibody HUTS21 (10 µg mL^−1^) + BDNF treated, and β1-integrin function blocking antibody 2999 (40 µg mL^−1^, applied immediately after injury) followed by Mn^2+^ + BDNF treated (applied 60 min after injury) zebrafish. All treatments were bath applied. Data are the mean ± standard error of the mean. (n>50, *P<0.01, ANOVA with Tukey’s post hoc analysis). BDNF: brain-derived neurotrophic factor; BSA: bovine serum albumin.

### Mammalian myelin factors block natural regeneration in the zebrafish spinal cord but are overcome by Mn^2+^ and BDNF treatment

Whereas zebrafish myelin breakdown products are permissive for axon outgrowth^26^, mammalian myelin factors suppress outgrowth of primary cultured zebrafish CNS neurons^26,30^. Does mammalian MAG attenuate natural regeneration in zebrafish? Delivery of mammalian MAG following zebrafish SCI robustly disrupted recovery of both the evoked C-start response and spontaneous swimming activity (Fig. 4a,b; Supplementary Fig. S7; Movie S4), but had no effect on activity when applied to uninjured fish (Supplementary Fig. S8). Administering BDNF together with Mn^2+^ (Fig. 4a,b; Supplementary Movie S5) or a β1-integrin activating antibody (Fig. 4b) after SCI and MAG exposure led to potent behavioral recovery. Importantly, BDNF or Mn^2+^ applied individually resulted in insignificant functional regeneration, as did BDNF in combination with Mg^2+^ or a non-activating β1-integrin antibody (Fig. 4b). Retrograde labeling of all crossing axon fibers, respectively, revealed that natural regenerative growth at the lesion site was strongly suppressed by MAG exposure (Fig. 4c). These effects were largely abolished with subsequent combined Mn^2+^ and BDNF treatment (Fig. 4c). Furthermore, administration of Mn^2+^ and BDNF together promoted robust M-cell re-growth even after initial MAG exposure (Fig. 4d). Altogether, these findings support the notion that mammalian MAG can inhibit zebrafish axon regeneration, which is reversed by a synergistic integrin activating and neurotrophin treatment. Experiments treating injured zebrafish with the mammalian CSPG aggrecan or Sema 3A also showed reduced levels of functional spontaneous swimming activity (Fig. 4e). Importantly, the activity of these inhibitory factors, which have been shown to function through negative regulation of β1-integrin, were also overcome by addition of Mn^2+^ and BDNF together (Fig. 4e).

**Figure 4.**
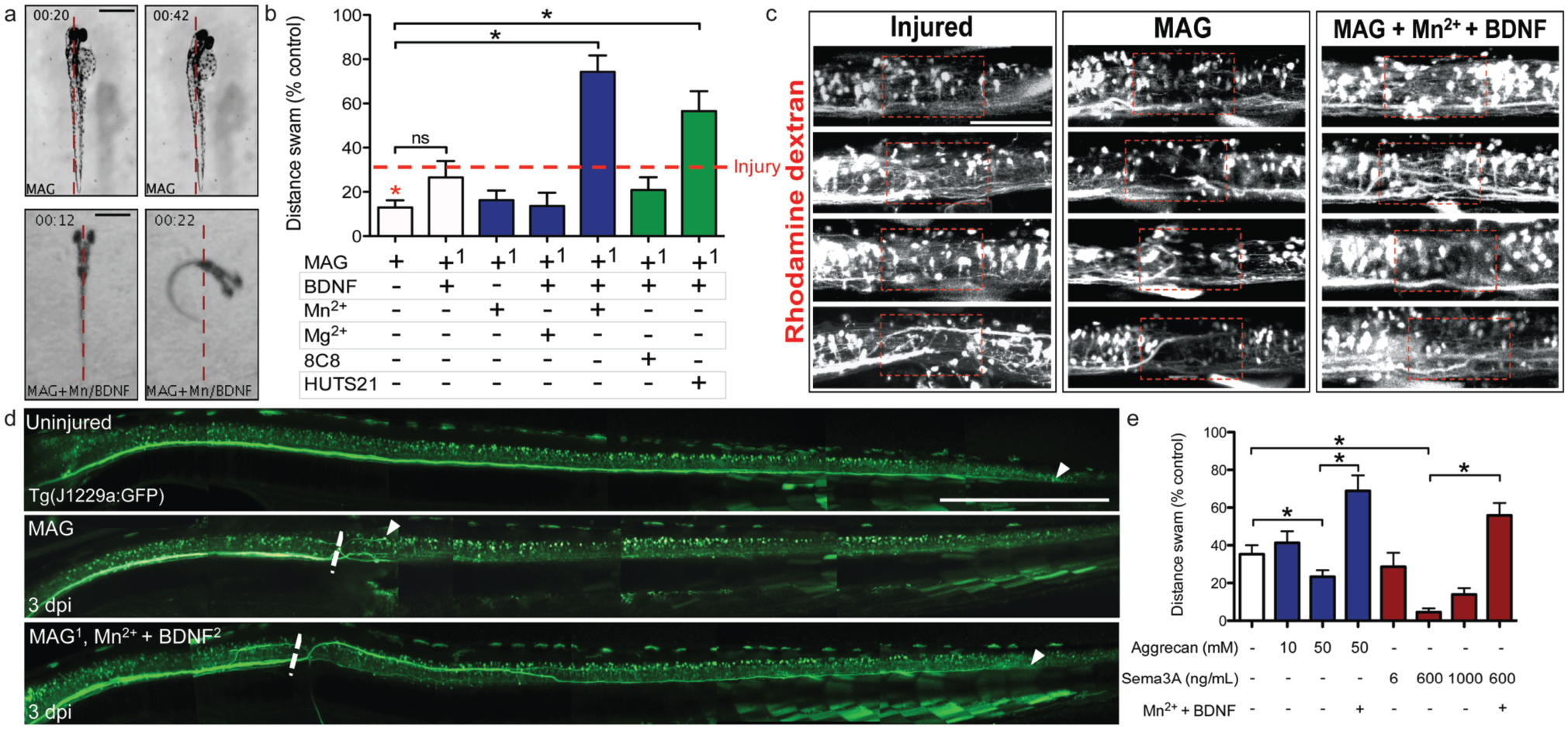
Mammalian myelin factors block natural regeneration in the zebrafish spinal cord but are overcome by Mn^2+^ and BDNF treatment. *a*, Evoked escape response “C-start” using high-speed video capture (1,000 Hz) depicting the frame immediately prior to response from a vibration stimulus and the frame demonstrating maximum body angle achieved for MAG treated (10 µg mL^−1^, applied immediately after injury) and MAG followed by administration of Mn^2+^ + BDNF (1 mM + 100 ng mL^−1^, 60 min after MAG treatment) treated zebrafish. Scale bar, MAG 1250 µm; MAG+Mn/BDNF 1,000 µm. *b*, Quantitative MATLAB analysis of spontaneous swimming activity of larval zebrafish at 3 d after complete spinal cord transection as a percentage of uninjured control-treated (BSA) zebrafish activity. Dashed line indicates the mean level of swimming behavior for injured control treated (BSA) zebrafish. *c*, Representative images from injured zebrafish spinal axons that were retrogradely labeled with tetramethylrhodamine dextran and were allowed to recover for 3 d with control-treatment (BSA), MAG (immediately after injury) or MAG (immediately after injury) and Mn^2+^ + BDNF treatment (60 min after MAG treatment). Red dashed box delineates original injury gap. Scale bar, 50 µm. *d*, Multi-photon imaging of the M-cell axon in the transgenic zebrafish line Tg(J1229a:GFP) of uninjured control treated (BSA), injured MAG treated, and injured MAG treated followed by Mn^2+^ + BDNF administration 60 min after injury to zebrafish. White arrowheads denote the most caudal region of the M-cell axon that was detected. Scale bar, 1 mm. *e*, Quantitation of spontaneous swimming activity for zebrafish treated with BSA (control), aggrecan (immediately after injury), Sema 3A (immediately after injury) or Mn^2+^ and BDNF (60 min after injury). Data are the mean ± standard error of the mean. BDNF: brain-derived neurotrophic factor; BSA: bovine serum albumin; MAG: myelin-associated glycoprotein.

## DISCUSSION

This study examined how targeting the functional state of integrins modulates the activities of MAG and BDNF on vertebrate spinal neurons. Our results indicate that stimulating integrin activity not only blocks negative remodeling of integrin receptors and inhibition of axon outgrowth by MAG, but also restores integrin clustering and stimulated outgrowth by BDNF. Moreover, targeting integrin activity noninvasively combined with BDNF treatment promoted regenerative growth of long projection axons in the zebrafish spinal cord that fail to naturally regrow after transection injury, leading to functional recovery. Exposure to mammalian MAG, CSPGs or Sema 3A blocked natural regeneration in the spinal cord, which was overcome by treatment to potentiate integrin activity combined with BDNF to restore motor function. Altogether, these findings indicate that the state of integrin activation plays complementary roles in modulating the output activity of opposing cues on integrin-based adhesions and supports functional nerve regeneration *in vivo*. Activators of integrin functional state may thus provide an effective therapeutic intervention to improve clinical outcomes.

The mechanisms underlying functional central nervous wiring and how those connections lead to complex, higher order cognitive functions remain some of the most challenging and elusive questions in modern biology. It has become clear that the answers will likely require a comprehensive understanding of how the molecular, cellular and systems-based levels interconnect. An important underpinning to understanding how a complex system functions is to comprehend its foundational infrastructure. Elucidation of the mechanisms used by pioneering growth cones may provide a basis for promoting restoration of nerve connections lost to injury or disease. Furthermore, several neurologic and psychiatric conditions have been associated with dysfunction in developmental wiring^30-32^. Thus, gaining a better appreciation for how these connections that are disrupted due to abnormalities in axon guidance affect normal function and lead to disease may offer insight into avenues for therapeutic intervention.

The complexity of the mammalian spinal cord milieu after injury creates a multitude of challenges both for axons attempting to regrow, and for investigations into therapies aimed at stimulating regeneration in the spinal cord, which to date have had limited success. Thus, a vertebrate animal model that possesses a relatively permissive environment after CNS injury may prove invaluable for elucidating methods to both enhance natural axon outgrowth, and trigger regeneration in neurons that demonstrate low intrinsic axon growth capacity after injury. The ability to couple this with the assessment of functional recovery utilizing the high-level innate predator avoidance response provides an additional advantage for this system. Our model expands on a previous report where delivery of cAMP into the M-cell soma after incomplete SCI promoted regenerative growth in the absence of exogenous inhibitory factors^22^. Whether our combined treatment paradigm may also elevate cAMP in the cell body, or whether elevated cAMP may regulate substrate adhesions are currently unknown, but warrant further investigation.

Neurotrophins, which play important physiologic roles in neuronal survival, axon outgrowth, and synaptic plasticity and maintenance throughout development and adulthood, make for attractive therapeutic candidates to promote regeneration after injury. Human clinical trials with systemic delivery of a neurotrophin showed little efficacy^34-36^. Current maxims suggest that these trials were unsuccessful for several reasons, including inability to efficiently penetrate the spinal cord lesion site, high side effect profiles, and possibly failure to promote axon outgrowth over local inhibitory factors such as MAG^14,37,38^. Thus, techniques for restoring neurotrophin effects on axon outgrowth may provide robust benefit for promoting regeneration in both the peripheral and central nervous systems.

Interestingly, with a single bath application of Mn^2+^ and BDNF treatment, we observed considerable regenerative growth of the Mauthner axon, which normally fails to demonstrate significant sprouting after injury. Whether Mn^2+^ and BDNF treatment has a direct impact on the Mauthner axon or cell body, or also affects support cells is unknown. Furthermore, how the Mauthner axon consistently elongates distally towards its previous synaptic destination, often finding the original axon tract even with occasional initial growth hundreds of microns outside of its stereotyped area, is under further investigation. Visualization of early Mauthner axon sprouting suggests that the preliminary response to transection is to grow in multiple available directions, often extending three or four individual axon sprouts. Is this seemingly successful targeting of the Mauthner axon an intrinsic response, whereby it re-expresses cell:to:cell transmembrane proteins that make growth optimal along its previous pathway supported by glia or fellow axon fibers? Or is there a molecular or cellular environmental response from either its synaptic target or local injury milieu that mediates appropriate guidance? Studies of peripheral motor axon regeneration suggest that regrowth commonly occurs along previous tracts, suggesting this appropriate regrowth response may be a conserved phenomenon^39^.

The present study demonstrates a new approach of targeting a downstream cellular process that mediates the functional effects of outgrowth inhibition while simultaneously acting to promote axon regenerative growth and robust functional recovery. Discovery of such strategies holds promise for promoting axon regeneration after traumatic injury, which is a critical step in restoring connectivity and functional recovery. Moreover, the novel strategy to add well-known barriers to mammalian regeneration exogenously to a relatively permissive vertebrate SCI model further illustrates and expands the utility of this approach.

## EXPERIMENTAL PROCEDURES

### Ethics statement

All animal experiments were carried out with adherence to National Institutes of Health (NIH) Guidelines for animal care and safety and were approved by the Mayo Clinic Institutional Animal Care and Use Committee (#A40811 and #A7912). Both sexes were used equally.

### Primary cell culture of Xenopus laevis and zebrafish spinal neurons

Experiments were conducted as described previously^14^. Briefly, primary spinal neurons were prepared from neural tube dissections of 1-d old (stage 22) *Xenopus laevis* embryos^40^. Both sexes were used equally. These cultures were used for experiments 10 h to 16 h after plating on fibronectin substrate at 20-22°C. All cover glasses were coated with fibronectin (20 µg mL^−1^, Sigma). *Xenopus* spinal neuron cultures were treated with mammalian MAG-Fc (1 µg mL^−1^, R&D Systems), MnCl_2_ (1 mM, Sigma), BDNF (50 ng mL^−1^, Peprotech) or a control BSA vehicle solution, followed by standard chemical fixation. MAG treatments for 5 min were utilized to obtain maximal integrin surface removal^6^. Zebrafish (*Danio rerio*) spinal cultures were conducted as described previously^30^. Briefly, zebrafish spinal neurons were prepared from high-pec stage (42 hpf) wild-type Segrest embryos. Both sexes were used equally. All experimental groups were assigned to randomized codes.

### Quantitative immunofluorescence microscopy

*Xenopus* and zebrafish spinal neuron cultures were chemically fixed in 4% paraformaldehyde and 0.01% glutaraldehyde for 20 min^6,14,30^. We immunolabeled un-permeabilized cells using a mouse monoclonal antibody to the extracellular domain of β1-integrin (8c8, 0.8 µg mL^−1^, University of Iowa Developmental Studies Hybridoma Bank) along with a rabbit polyclonal anti-β-tubulin antibody (0.4 µg mL^−1^, Abcam ab15568, Cambridge, England) to exclude permeabilized cells. Zebrafish neurons were stained for activated β1-integrin (HUTS21; 1 mg mL^−1^; gift of the late Richard Pagano) and phospho-FAK (FAK^397^, ABT135; 0.8 mg mL^−1^; Millipore). Alexa dye-labeled secondary antibody conjugates (Invitrogen, Carlsbad, CA, USA) were used at 2 µg mL^−1^. Image analysis and processing was conducted as described previously^14^. Briefly, to measure only receptors at the plasma membrane, permeabilized growth cones were excluded from the analysis, as identified by tubulin immunofluorescence. The original 14-bit images were analyzed using ImageJ (Bio-Formats ZVI plug-in, Madison, WI, USA). Data were background subtracted and normalized to the appropriate control images. For quantification of integrin clustering, we used a three-fold fluorescence inclusion criterion of β1-integrin cluster intensity over the mean background fluorescence in the growth cone central domain as validated previously^14^. Analysis of primary spinal neuron outgrowth rates was conducted as previously described^14^. In short, cultured spinal neurons were recorded immediately before treatment intervention and followed for at least 60 min (ProgRes CapturePro 2.7, Jenoptik Inc., Jupiter, FL, USA). All outgrowth assays were performed on a Zeiss 40 compact fluorescent lamp microscope equipped with a Ludl Electronic Products (Hawthorne, NY, USA) BioPoint 2 motorized stage, cooled charged-coupled device camera and a 20X objective.

### Larval zebrafish maintenance and spinal cord injury

Zebrafish embryos were obtained (<15 h post fertilization) from the Mayo Zebrafish Core facility and maintained in petri dishes containing zebrafish E3 embryo water in a 28°C incubator (12 h:12 h light:dark cycle). Both sexes were used equally. Larval zebrafish (3 dpf) were anesthetized by adding 0.17% tricaine into the zebrafish embryo water for 1 min. Anesthetized zebrafish were then transferred by using a fire-polished Pasteur pipette to a silica grooved chamber (groove diameter, 5-10 mm) filled with 0.17% tricaine zebrafish embryo water to maintain anesthesia. A fine pulled glass micro-needle was used under control of a stereomicroscope to make a complete spinal cord transection at the pre-caudal vertebral level (myotome 7-9). After injury, larval fish were immediately transferred to a petri dish filled with embryo water and maintained in a 28°C incubator and monitored closely during a 1-h recovery period. Once in the recovery dish, but before motor control was regained, fish were randomly transferred to coded experimental dishes. Nonresponsive fish, fish that were bent along their body axis, or fish that did not regain fin movement were removed from the dish and euthanized. Euthanasia was consistent with the recommendations of the 2013 AVMA Guidelines on Euthanasia. Double-blinded quality control analysis using two-photon imaging after injury validated the consistency of complete spinal cord transection in larval zebrafish (Supplementary Fig. S3). All treatments were bath administered 60 min post-injury (BDNF 100 ng mL^−1^; Mn^2+^ 1 mM; HUTS21 10 mg mL^−1^), except for MAG (10 mg mL^−1^) and the β1-integrin function-blocking antibody (2999; 40 mg mL^−1^; gift of Kenneth Yamada, NIH), Sema3A-Fc (R&D Systems), and the CSPG aggrecan (Sigma), which were added immediately after transection.

### Downregulating β1-integrin expression

Lissamine-tagged antisense morpholino oligonucleotides were characterized previously to target zebrafish β1-integrin co-orthologs (itgb1a 5’-TATGAAAAGTAGCTTCAGGTCCATC-3’, itgb1b-1 5’-GGAGCAGCCTTACGTCCATCTTAAC-3’, itgb1b-2 5’-GCCAGTTTGAGTGAATAACTCACCT-3’; Gene Tools, LLC)^29^. A scrambled, non-targeting lissamine-tagged morpholino was used as a control (scMO 5’-CCTCTTACCTCAGTTACAATTTATA-3’). Morpholino knockdown of the β1-integrin co-orthologs was conducted by local injection (1-2 mg mL^−1^) using a pulled micro-needle under standard pulse pressure controlled by a Picospritzer delivered immediately following transection injury.

### Zebrafish behavioral testing

The evoked predatory escapes response, or C-start, was adapted from McGee and colleagues^41^. Briefly, a trigger-activated system connected to an electronic chip delivered a short vibrational stimulus to the zebrafish chamber. The C-start response was recorded using a high-speed digital video camera (Redlake MotionScope M1, Tucson, AZ) at 1,000 Hz, which was mounted to a Zeiss stereomicroscope. Movie files were analyzed using ImageJ software, using randomly assigned codes for the experimental groups. During zebrafish activity assays, larval zebrafish were placed individually into a single well of a 24-well plate and set on top of a lighted LED-platform. Zebrafish were allowed to acclimate to a sound-controlled room for at least 30 min. Activity was then recorded for 60 min using a mounted video camera above the chambers. Assessment of the recorded activity was automated using a MATLAB GUI program (written by David Argue, Mayo Clinic).

### Zebrafish whole-mount and live imaging

Zebrafish used for whole-mount imaging were chemically fixed in 4% paraformaldehyde and 4% sucrose at room temperature for 3 h, then permeabilized with Proteinase K (50 mg mL^−1^) for 30 min. Zebrafish were then placed in blocking buffer (10% normal goal serum, 1% DMSO, 0.8% Triton-X100) for 1 h at room temperature. Embryos were incubated in primary antibody (3A10, 1:20; znp-1, 1:50; University of Iowa Developmental Studies Hybridoma Bank) overnight at 4°C, followed by incubation with secondary antibody (Alexa555 A-21147, Alexa488 A-11029; 1:500, Invitrogen) overnight at 4°C. For retrograde labeling, zebrafish were injured and received a local injection at the lesion site of tetramethylrhodamine dextran (10,000 MW, Invitrogen, D1817). During live imaging, zebrafish were anesthetized by adding 0.17% tricaine into the zebrafish embryo water for 1 min. Both whole-mount immunostained and live zebrafish were mounted in clear tubing (Fisher, #05-701-41) and imaged using a multi-photon microscope (FV1,000 multi-photon excitation (MPE) microscope, Olympus). All zebrafish experimental groups were assigned random codes for double-blinded imaging and behavioral testing.

### Statistical analyses

The sample size for all experiments was based on a power analysis using a one-way ANOVA with Tukey’s post hoc analysis, a 0.05 significance value, and achieving >80% power. All statistical analyses were performed using GraphPad Prism software (v5, La Jolla, CA, USA). The figure legends state the statistical tests used. Data with a normal distribution (D’Agostino and Pearson omnibus normality test) were assessed using repeated-measures one-way analysis of variance with Tukey’s post hoc analysis. Variation within each group of data are reported as either s.d. or s.e.m., as indicated in the text and figure legends. Statistical comparisons were made between groups of data with similar variance.

## Acknowledgements

Funding by the National Institutes of Health (NS67311, JRH; T32 GM065841, LPC; NS080322, TRC), a John M. Nasseff, Sr. Career Development Award in Neurologic Surgery Research from the Mayo Clinic (JRH), and a Langan M.D., Ph.D. fellowship (LPC) supported this work. The content is solely the responsibility of the authors and does not necessarily represent the official views of the National Institutes of Health. We appreciate the Mayo Microscopy and Cell Analysis Core for experimental and technical support. We would like to thank David Argue (Mayo Clinic) for customization of the MATLAB GUI program for quantitating swimming activity, Tammy Greenwood (Mayo Clinic) for assistance setting up and maintaining fish in the Mayo Clinic Zebrafish Core Facility, Fredric Meyer (Mayo Clinic) for stimulating discussions, Hitoshi Okamoto (RIKEN Brain Institute, Wako, Japan) for the *Tg(J1229a:GFP)* transgenic line, Ken Yamada (US National Institutes of Health) for the 2999 β1-integrin function blocking antibody and helpful discussions, the late Richard Pagano for the HUTS21 antibody, Ben Meyer, Jarred Nesbit and the entire Henley lab for support and technical assistance.

## Author contributions

LPC, JRH, TRC, HLS, MRM, and SCE devised experiments; LPC and TRC performed experiments; LPC, TRC and JRH analyzed data; LPC and JRH wrote the manuscript. All authors read and provided critical feedback on the manuscript.

## SUPPORTING INFORMATION (See Aditional Files)

**Supplementary Figure S1. β1-integrin activating antibody and BDNF overcome growth inhibition by MAG in cultured *Xenopus* spinal neurons.** Quantitation of live rates of axon outgrowth from primary *Xenopus* spinal neurons treated with β1-integrin activating antibody, HUTS21, BDNF and HUTS21, MAG, or MAG followed by HUTS21 + BDNF. Colored dash lines represent the mean of individual application of MAG followed by Mn^2+^ and BDNF treatment (green), and Mn^2+^ alone (red). Data are the mean ± standard error of the mean. (n > 100, *P < 0.05, ANOVA with Tukey’s post hoc analysis).

**Supplementary Figure 2. Quality control analysis of complete spinal cord transection procedure in larval zebrafish.** *a*, Phase image of rostral half of larval zebrafish, 3 dpf; red dashed box delineates the target region of spinal cord injury. *b*, Example images of uninjured and injured Tg(olig2:EGFP) zebrafish^42^; all images from blinded quality control injury trial. Scale bar, 50 µm. *c*, Example images of uninjured and injured Tg(J1229a:GFP) zebrafish; all images from blinded injury trial. Scale bar, 50 µm.

**Supplementary Figure S3. Activation of zebrafish primary spinal neuron β1-integrins by Mn^2+^.** *a*, Representative images of zebrafish primary spinal neurons treated with BSA (control) or Mn^2+^ and stained for activated β1-integrins or phosphorylated FAK^397^. *b*, Quantification of fluorescence intensity from BSA (control) or Mn^2+^ treated neurons. Data are the mean ± standard error of the mean. *c*, Validation of β1-integrin antibody using morpholino knockdown. Surface expression of N-cadherin receptor was unaffected by β1-integrin morpholinos.

**Supplementary Figure S4. Regenerative growth induced by BDNF after complete SCI.** Larval zebrafish were treated with BSA (injured) or BDNF and final M-cell axon regenerative growth was recorded 3 dpi using whole mount immunofluorescence (3A10). Data are the mean ± standard error of the mean. (*P<0.05, ANOVA with Tukey’s post hoc analysis).

**Supplementary Figure S5. Functional recovery through targeting integrin receptors.** Evoked escape response “C-start” assay depicting high-speed video capture (1,000 Hz) of larval zebrafish from time of vibration stimulus delivery to first recorded movement (*a*), and duration of movement to reach initial maximal body axis bend (*b*) for control-treated (BSA), Mn^2+^ or BDNF individually treated and combined Mn^2+^ + BDNF treated injured zebrafish. Quantification as a percentage of control uninjured at one and 3 d after injury. Data are the mean ± standard error of the mean. (n>15, *P<0.05, ANOVA with Tukey’s post hoc analysis).

**Supplementary Figure S6. Downregulating β1-integrin expression disrupts natural regeneration and abolishes regeneration induced by Mn^2+^ and BDNF.** Swimming activity assay for injured zebrafish that received targeted delivery with a control non-targeting morpholino, or two morpholino combinations to knockdown both β1-integrin co-orthologs followed by bath treatment with BSA or with Mn^2+^ and BDNF. Data are the mean ± standard error of the mean. (n>15, *P<0.05, ANOVA with Tukey’s post hoc analysis).

**Supplementary Figure S7. MAG disrupts zebrafish C-start response.** Frequency of elicited C-start response to a vibration stimulus for uninjured control treated (BSA), injured control treated (BSA), MAG treated (immediately after injury), or MAG (immediately after injury) and Mn^2+^ + BDNF treated (60 min after MAG treatment) zebrafish. Data are the mean ± standard error of the mean. (n>15, *P<0.05, ANOVA with Tukey’s post hoc analysis).

**Supplementary Figure S8. No impairment of uninjured zebrafish swimming activity by MAG.** Swimming activity for uninjured zebrafish (3 dpf) that received control treatment (BSA) or MAG recorded at 1 and 3 d after administration. Data are the mean ± standard error of the mean. (n>25, *P<0.05, ANOVA with Tukey’s post hoc analysis).

**Supplementary Movie S1.** Evoked escape response “C-start” assay depicting high-speed video capture (1,000 Hz) of larval zebrafish, treated with BSA. Vibration stimulus delivered at time point 00:00. The video corresponds to Fig. 3A control.

**Supplementary Movie S2.** Evoked escape response “C-start” assay depicting high-speed video capture (1,000 Hz) of larval zebrafish 3 d after complete transection, treated with BSA. Vibration stimulus delivered at time point 00:00. The video corresponds to Fig. 3a injury.

**Supplementary Movie S3.** Evoked escape response “C-start” assay depicting high-speed video capture (1,000 Hz) of larval zebrafish 3 d after complete transection, treated with Mn^2+^ and BDNF (60 min after injury). Vibration stimulus delivered at time point 00:00. The video corresponds to Fig. 3a Mn^2+^ and BDNF treatment after injury.

**Supplementary Movie S4.** Evoked escape response “C-start” assay depicting high-speed video capture (1,000 Hz) of larval zebrafish 3 d after complete transection, treated with MAG (immediately after injury). Vibration stimulus delivered at time point 00:00. The video corresponds to Fig. 4a MAG treatment after injury.

**Supplementary Movie S5.** Evoked escape response “C-start” assay depicting high-speed video capture (1,000 Hz) of larval zebrafish 3 d after complete transection, treated with MAG (immediately after injury) and Mn^2+^ + BDNF (60 min after MAG treatment). Vibration stimulus delivered at time point 00:00. The video corresponds to Fig. 4a MAG plus Mn^2+^ and BDNF treatment after injury.

